# Genome-wide transcriptional profiling uncovers a similar oligodendrocyte-related transcriptional response to acute and chronic alcohol drinking in the amygdala

**DOI:** 10.1101/2021.09.07.459347

**Authors:** Sharvari Narendra, Claudia Klengel, Bilal Hamzeh, Drasti Patel, Joy Otten, Roy Lardenoije, Emily L. Newman, Klaus A. Miczek, Torsten Klengel, Kerry J. Ressler, Junghyup Suh

## Abstract

Alcohol intake progressively increases after prolonged consumption of alcohol, but relatively few new therapeutics targeting development of alcohol use disorder (AUD) have been validated. Here, we conducted a genome-wide RNA-sequencing (RNA-seq) analysis in mice exposed to different modes (acute vs chronic) of ethanol drinking. We focused on transcriptional profiles in the amygdala including the central and basolateral subnuclei, a brain area previously implicated in alcohol drinking and seeking, demonstrating distinct gene expression patterns and canonical pathways induced by both acute and chronic intake. Surprisingly, both drinking modes triggered similar transcriptional changes, including up-regulation of ribosome-related/translational pathways and myelination pathways, and down-regulation of chromatin binding and histone modification. Notably, multiple genes that were significantly regulated in mouse amygdala with alcohol drinking, including *Atp2b1, Slc4a7, Nfkb1, Nts*, and *Hdac2*, among others had previously been associated with human AUD via GWAS or other genomic studies. In addition, analyses of hub genes and upstream regulatory pathways predicted that voluntary ethanol consumption affects epigenetic changes via histone deacetylation pathways, oligodendrocyte and myelin function, and oligodendrocyte-related transcriptional factor, *Sox17*.

Overall, our results suggest that the transcriptional landscape in the central and basolateral subnuclei of the amygdala is sensitive to voluntary alcohol drinking. They provide a unique resource of gene expression data for future translational studies examining transcriptional mechanisms underlying the development of AUD due to alcohol consumption.

## Introduction

Alcohol use disorder (AUD) is a chronic relapsing brain disorder that is a major public health concern in the United States, where the lifetime prevalence of AUD among adults is nearly 30% [1]. Despite the disorder’s prevalence and severity, there are few effective treatments for alcohol abuse. The hallmark of AUD is gradually increasing alcohol consumption over time [2]. This increase in alcohol intake is thought to result from neurobiological adaptation induced by alcohol [3]. Prolonged heavy alcohol exposure appears to cause progressive dysfunction in multiple brain areas, most notably changes in neuronal plasticity in the brain’s reward and stress systems such as the amygdala [4].

The amygdala is comprised of multiple interconnected nuclei nested deep in the temporal lobe in human and its structures and functions are well-conserved across evolution. It has been associated with both emotion and motivation, playing an essential part in processing aversive and appetitive information [5-7]. Previous neuroimaging studies demonstrated that alcohol cues trigger amygdala activation which correlates with craving for alcohol in patients with AUD [8, 9]. In animal models, chronic alcohol exposure alters neuronal transmission in the central nucleus of the amygdala (CeA) and the neural activity of the CeA during alcohol withdrawal is associated with levels of alcohol drinking in alcohol-dependent rats [10, 11]. Furthermore, the activation of the basolateral amygdala (BLA) and its projections to the nucleus accumbens is necessary for cue-induced alcohol seeking behaviors [12].

As alcohol has multiple direct molecular targets, identifying and characterizing genes in a brain region-specific manner is vital to our understanding of the molecular mechanisms underlying alcohol-related behaviors and AUD development and susceptibility [13-16]. Several studies have applied genomics to examine alcohol-induced transcriptional effects using chronic models of voluntary ethanol consumption and forced vapor exposure through ethanol vapor in rodents [17-21]. The results identified multiple molecular targets, such as alterations in neuronal function and signal transduction, indicating that chronic ethanol exposure and withdrawal have prominent actions on gene expression in multiple brain areas including the prefrontal cortex. However, these studies using microarrays with pre-determined numbers of genes and forced alcohol exposure have not directly addressed genome-wide transcriptional response to voluntary alcohol drinking. In addition, few studies investigated gene networks that are targeted by voluntary alcohol drinking in the amygdala where molecular processes may underlie the development and maintenance of alcohol-drinking and seeking behaviors [22, 23].

To acquire a better insight into gene expression alterations impacted by acute and chronic voluntary oral ethanol consumption, we employed a 2-bottle choice ethanol drinking procedure with either a single bout or chronic intermittent access that has been shown to escalate ethanol intake over weeks in female mice [24]. We then explored transcriptomic changes in the amygdala that may underly progressive increase in ethanol intake. We found that acute and chronic ethanol drinking induced similar network-level changes in gene expression, suggesting that a single episode of ethanol consumption substantially alters amygdala transcriptomes that may last for a long time. Furthermore, we identified expression networks that correlated with the level of ethanol consumption and ethanol preference, suggesting mechanistic relationships. Our bioinformatics analyses also revealed that some of the most strongly correlated genes include myelination, synaptic transmission, chromatic modification, translation, and RNA processing. Together, our findings provide systems-level evidence of the relationships between voluntary alcohol drinking and gene networks within the central and basolateral subnuclei of the amygdala.

## Materials and Methods

### Animals

Two separate cohorts (N = 12 for the first and N = 18 for the second cohorts) of adult female C57Bl/6J mice at 7 weeks of age were purchased from Jackson Laboratories (Bar Harbor, ME) and kept under standard conditions with 12:12-hour light/dark cycle (lights on: 07:00). Animals were group housed upon arrival and acclimated for 1-2 weeks. Then, mice were individually housed and allowed access to tap water and free (ad libitum) access to standard laboratory chow during the whole experimental period. All experiments were approved by and carried out in accordance with the Institutional Animal Care and Use Committee at McLean Hospital. All experimental and animal care procedures met guidelines outlined in the NIH Guide for the Care and Use of Laboratory Animals. All efforts were made to minimize distress and the number of animals used.

### Alcohol drinking procedures

Twenty percent of ethanol solution (v/v) was prepared in tap water from 95% ethyl alcohol (Pharmaco-AAPER, Brookfield, CT). Mice were changed to individual housing at least 24 hours before the presentation of 2 plastic tubes of water on the cage lid for 2 days for acclimation to drinking from sipper tubes. Fluids were presented in 50-ml conical tubes (Falcon) with no. 6 rubber stoppers (#6R, Ancare, Bellmore, NY) containing stainless steel ball-bearing sippers (TD-100, Ancare). Centrifuge tubes were securely held through the metal wire cage lid and presented to mice 2 hours before the dark cycle. Bottles were weighed to the nearest hundredth of a gram 24 hours after the fluids were given and the left/right position of the bottles were alternated before each ethanol drinking session to avoid side preferences. To control for spillage and evaporation, daily “drip” averages (loss of fluid in two cages with no animal present) were subtracted from individual fluid intakes. Mice were also weighed weekly to the nearest tenth of a gram to calculate the grams of ethanol intake per kilogram of body weight. Preference for ethanol was calculated for ethanol compared with water, with formula being volume of ethanol intake (ml) divided by total volume fluid intake (ml).

Mice from each cohort were assigned to three drinking groups. Mice in the acute drinking group (Acute Drinking) were given two bottles of water for 27 days, then a bottle with 20% ethanol and a bottle with water on Day 28. The chronic intermittent access drinking group (Chronic Drinking) of mice received free-choice 24-hour access to 20% ethanol and water on every-other-day (EOD) basis for 4 weeks (28 days). Mice in the water drinking group (Water Drinking) received the same schedule of total fluid access but consumed only water from two bottles. After completion of the experiments on day 29, at 1-2 hours after lights on, mice were deeply anesthetized with isoflurane and sacrificed by decapitation. Trunk blood was collected in EDTA tubes to measure blood ethanol concentration (BEC), and vaginal smear was collected on non-coated glass microscope slides to determine the stage of the estrous cycle. Brains were rapidly removed from skull, and placed on dry ice, and stored at -80 °C until further processing. Fresh frozen brains were sectioned at a thickness of 300 um, then micropunches (1 mm in diameter and 1 mm in thickness) were bilaterally collected from the entire amygdala including basolateral amygdala (BLA) and central amygdala (CeA) based on established anatomical coordinates from the mouse brain atlas [25]. The micropunches were aimed to include the following coordinates: ML ±3.2, AP -1.5, DV -5.0 mm, and all the samples were placed in microcentrifuge tubes (1.5 ml), kept frozen in dry ice, and stored at -80 C until RNA isolation. BEC was determined using the Analox Analyzer (Analox Instruments Inc., Lunenburg, MA) from blood samples (30 ul). The vaginal cytology was carried out using crystal violet staining [26].

### RNA extraction and sequencing

Total RNA was isolated and purified using the Absolutely RNA Miniprep Kit (Cat# 400800, Agilent Technologies, Santa Clara, CA, USA) according to the manufacturer’s protocol for extremely small samples. Quality and concentration of the extracted RNA were evaluated using a NanoDrop 8000 spectrophotometer (Thermo Scientific). Eleven samples from the first batch (4 from Water, 4 from Acute and 3 from Chronic Drinking groups) and 17 samples from the second batch (6 from Water, 5 from Acute and 6 from Chronic Drinking groups) were sent to Beijing Genomics Institute (Hong Kong, China). Library construction and whole genome sequencing were conducted on BGISEQ-500 platform using the DNBseq short-read 100 bp paired-end reading.

### Data Processing

Raw sequencing reads were quality assessed with FastQC (https://www.bioinformatics.babraham.ac.uk/projects/fastqc/) (Babraham Institute, Cambridge, UK). Salmon [27] was used to first build a transcriptome index using the reference transcriptome for *Mus musculus* (GRCm38/mm10) along with the non-coding RNAs of the same, and then to quantify the RNA-seq samples at the transcript level. The R package, tximport, [28] was used to import this transcript-level abundance generated by Salmon, the estimated counts and the corresponding transcript lengths, and summarize it into transcript-level and gene-level expression matrices. Gene-level expression matrix was generated using the GENCODE mouse annotation (www.gencodegenes.org) as reference (release M21).

### Estimation of cell type abundances

Cell type abundances were estimated with BRETIGEA, an R package [29], for each of the following cell types – neuron, oligodendrocyte, microglia, oligodendrocyte precursor cell, astrocytes, and endothelial cell. BRETIGEA contains thoroughly validated datasets containing brain cell type-specific marker genes. Using the ‘*brainCells’* function, with the filtered dataset containing 16,260 genes as input, a sample-by-cell type matrix of estimated cell type proportion variables were generated.

### Differential expression

ComBat-seq [30] was used to adjust the batch effect by keeping the negative binomial distribution of RNA-seq reads count and the integer nature of the data. A pre-filtering step was then applied to the corrected gene counts where genes with less than ten counts on average across all the samples were filtered out, after which 16,260 genes were retained for the subsequent analysis. The R package DESeq2 [31] was used for downstream differential expression analysis using default parameters. The readings were then normalized using DESeq2 with the means of normalized counts as a filter statistic. In the initial data exploration, principal component analysis (PCA) was constructed to calculate the coefficient of variation between groups directly using ‘*plotPCA’*, a functionality in DESeq2.

### Functional and Pathway Enrichment Analysis of DEGs

The lists of DEGs were first divided into two sets, up- and down-regulated, with the criteria of p-value cutoff < 0.05, which was set to increase the number of genes in each condition. Then, function and pathway enrichment analysis were performed using the R package, clusterProfiler [32]. Gene ontology (GO) analysis provided gene annotations in biological processes (BP), molecular functions (MF), and cellular components (CC). In addition, Kyoto Encyclopedia of Genes and Genomes (KEGG) analysis provided more information on biological pathways related to diseases and drug targets.

### GWAS Catalog and DisGeNET analysis

To obtain a better understanding of whether the significant genes obtained in this study were already implicated in previously published alcohol addiction studies, the list of significant genes from the DESeq2 analysis was compared with the list of significant genes in GWAS Catalog (www.ebi.ac.uk/gwas) and DisGeNET (www.disgenet.org) datasets. The “*All associations v1*.*0*.*2 -with added ontology annotations, GWAS Catalog study accession numbers and genotyping technology*” dataset from the GWAS Catalog website and the “*ALL gene-disease associations*” dataset from the DisGeNET website were first downloaded and analyzed in R. From the GWAS Catalog dataset, the columns of interest “*DISEASE/TRAIT*” and “*MAPPED_GENE*” were retained, while from the DisGeNET dataset, “*geneSymbol*” and “*diseaseName*” were retained. The analysis was performed with disease terms containing “*alcohol*” from both “*DISEASE/TRAIT*” and “*diseaseName*” columns, after which, the disease terms containing “*nonalcohol/non-alcohol*” were filtered out. The significant genes from the DESeq2 analysis were then matched with the resulting genes from both the GWAS Catalog and DisGeNET datasets, to get the overlapping/common genes.

### Identification of Hub genes and Regulatory Transcription Factors

To visualize protein-protein interactions networks among DEGs, the Search Tool for the Retrieval of Interacting Genes (STRING) online database (version 11.0) [33] was used with DEGs with the criteria of adjusted p-value cutoff < 0.05. The same sets of DEGs were also used with GeneGo MetaCore (Clarivate Analytics, PA, USA) to detect upstream transcription factors. The Transcriptional Regulatory Relationships Unraveled by Sentence-based Text mining (TRRUST) (version 2) [34] online database was also utilized to discover transcriptional regulatory networks.

### cDNA synthesis and quantitative PCR (qPCR)

RNA samples were reverse transcribed into cDNA using superscript IV kit (cat# 18091200, Thermo Scientific, Waltham, MA) using random hexamer primers. Complementary DNA was amplified on a ViiA7 Real-Time PCR system (Thermo Scientific) with POWRUP SYBR Green Master Mix (cat# 4368706Thermo Scientific). Primers for genes of interest and housekeeping gene *Actb* were as follows: *Bc1* (fwd. 5’ GGTCCTCAGCTCTGGAAAAA 3’; rev. 5’ AGGTTGTGTGTGCCAGTTACC 3’), *Btg2* (fwd. 5’ GCGAGCAGAGACTCAAGGTT 3’; rev. 5’ CCTTTGGATGGTTTTTCTGG 3’), *Haghl* (fwd. 5’ GGACTCACCAGCCCTCTTCT 3’; rev. 5’ TTGGCCAAGCTCTGGTACAT 3’), *Kdm3a* (fwd. 5’ CATTGGAGCAAAACTTCCTCA 3’; rev. 5’ TGGTTTTGTTCTCGGTACTTCA 3’), *Lrrc24* (fwd. 5’ GCTGGATTTCACCTTCTTGC 3’; rev. 5’ GCCTGGTCCTCCAGTAATTC 3’), *Nenf* (fwd. 5’ GGATCCAGCAGACCTCACTC 3’; rev. 5’ TGGCTTTGTACACCTTGCTG 3’), *Nts* (fwd. 5’TCCAGCTCCAGAAAATCTGC 3’; rev. 5’CCTTCTCGTTTTTATCATTGACG 3’), and *Actb* (fwd. 5’ CCAACCGTGAAAAGATGACC 3’; rev. 5’ACCAGAGGCATACAGGGACA 3’). Specificity of the qPCR reaction was confirmed with melt curve analysis to ensure that only the expected PCR product was amplified. Duplicates were run for each reaction, and Ct values were normalized using the established delta-delta Ct method (2–ΔΔCt) and then normalized to *Actb* Cts.

### Statistical analysis

Data were analyzed using R or Graphpad Prism (version 9.1, GraphPad Software, San Diego, CA). The level of significance was set at p ≤ 0.05, and results are presented as mean plus or minus standard error of the mean (*M* ± SEM). For the drinking data, ethanol intake (g/kg), volume (ml) of water and ethanol consumed, total fluid intake (ml), ethanol preference (%), and body weight (g) were analyzed with multiple two-way analyses of variance (ANOVAs), followed by Bonferroni post hoc analysis when significant group effects were found (p < 0.05). BEC (mg/dl) and single daily ethanol intake (g/kg) on day 29 between Acute and Chronic Drinking groups were analyzed with one-way ANOVA.

## Results

### 2-bottle choice drinking

We employed a well-established 2-bottle choice drinking paradigm [24] and divided mice into 3 drinking groups (Water, Acute and Chronic EtOH) (Figure 1a). Ethanol intake of the first cohort was slightly higher than that of the second cohort in both Acute and Chronic Drinking groups but the small difference was not statistically significant. Consistent with previous behavioral studies [24, 35], mice in the Chronic Drinking group increased ethanol intake across the first 2-week period, subsequently maintaining a stable level (a daily average of 23.38 ± 0.78 g/kg in weeks 3-4). On day 28, mean ethanol intake was 20.96 ± 2.00 g/kg for the Chronic Drinking group, which was significantly higher (p < 0.05) than that of the Acute Drinking group (11.41 ± 1.54 g/kg) (Figure 1b). As expected, there were no differences in total liquid consumption across 4 weeks among the groups (Figure 1c). Consequently, analysis of 24-hour preference values revealed that mice in the Chronic Drinking group showed increased preference as early as after 2 weeks of intermittent drinking, which was sustained at an average 72.60 % in weeks 3-4. In contrast, mice in the Acute Drinking group displayed 45.98 % preference on Day 28 (Figure 1d). During 4 weeks of the drinking period, body weight (g) did not show any significant group differences on day 1 (Water, 19.60 ± 0.51; Acute, 19.63 ± 0.36; Chronic, 19.71 ± 0.32) and day 29 (Water, 20.83 ± 0.47; Acute, 20.54 ± 0.23; Chronic, 20.81 ± 0.41). On the final day of the study, all the mice in each drinking group were in either proestrus or estrus phase of the estrous cycle. There is no difference in alcohol consumption between mice in different phases, which is consistent with previous findings that alcohol intake is not affected by estrous cycle phase in female rodents [36, 37]. These data demonstrate that our protocol succeeded in achieving standard levels of both acute and chronic drinking behaviors and metabolism for subsequent transcriptional profiling of amygdala function.

**Figure 1.**
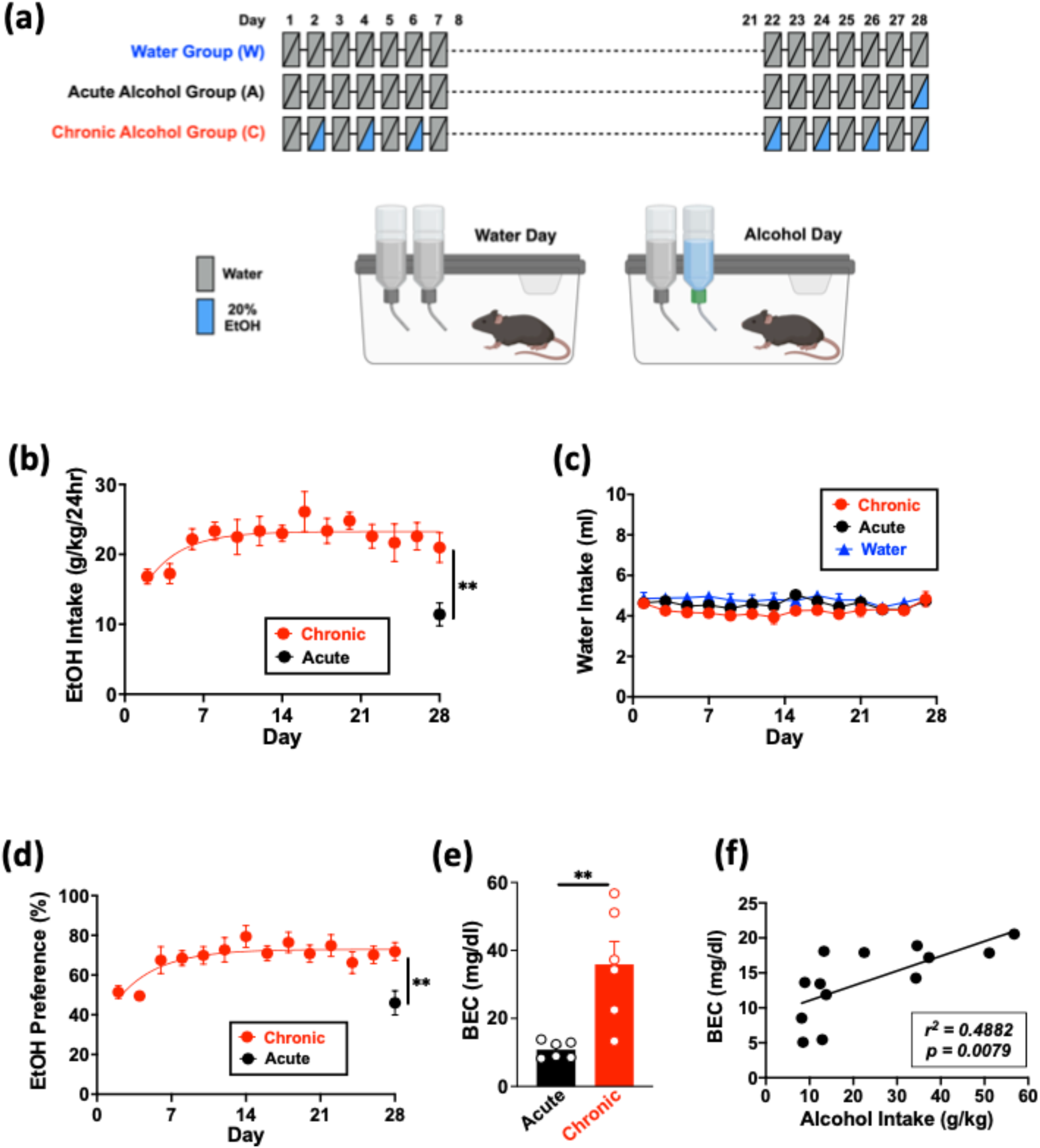
Experimental design and fluid consumption levels. (**a**) Experimental design and timeline. (**b**) Ethanol intake (g/kg BW) over 24 hours on water/20% EtOH drinking days. (**c**) Water intake over 24 hours on water/water drinking days. (**d**) EtOH preference ratios. (**e**) Blood ethanol concentration (mg/dl) measured in Acute and Chronic Drinking groups of 2nd cohort of mice on Day 28. (**f**) Correlation of BEC to the amount of ethanol consumed by mice in Acute and Chronic Drinking groups on Day 28. Data are mean ± SEM. ** *p* < 0.01 difference between groups.

### RNA-seq analysis

Since acute and chronic excessive alcohol consumption leads to gene expression alterations and cellular adaptations, RNA sequencing (RNA-seq) analysis was used to determine genome-wide transcriptomic profiles. Given the important roles of central amygdala (CeA) and basolateral amygdala (BLA) in alcohol-related synaptic changes and behaviors, we collected micropunches containing these subnuclei from three Drinking groups: Water, Acute and Chronic EtOH (Figure 2a). Since we collected RNA samples from two independent cohorts of mice, we first used Combat-seq [30] to adjust for any batch effects by keeping the negative binomial distribution of RNA-seq reads count and the integer nature of the data. Then, among 16,260 genes detected in the results, we assessed overall similarity among the samples by utilizing an unsupervised classification method, principal component analysis (PCA), and confirmed no outliers isolated by experimental conditions along two first principal components with 55% of the total variance (Figure 2b). Furthermore, since differences in cell type proportions can be a major source of variation in gene expression profiles, we used a computational cell type deconvolution tool, BRETIGEA [29] to estimate the abundances of six relevant cell types, including neurons, astrocytes, microglia, oligodendrocyte precursor cells (OPC), oligodendrocytes, and endothelial cells. We found no significant differences in any of the assessed cell types (Figure 2c). Together, these findings suggest that most of the observed variation in gene expression can be attributed to molecular implications of alcohol consumption rather than other confounding factors.

**Figure 2.**
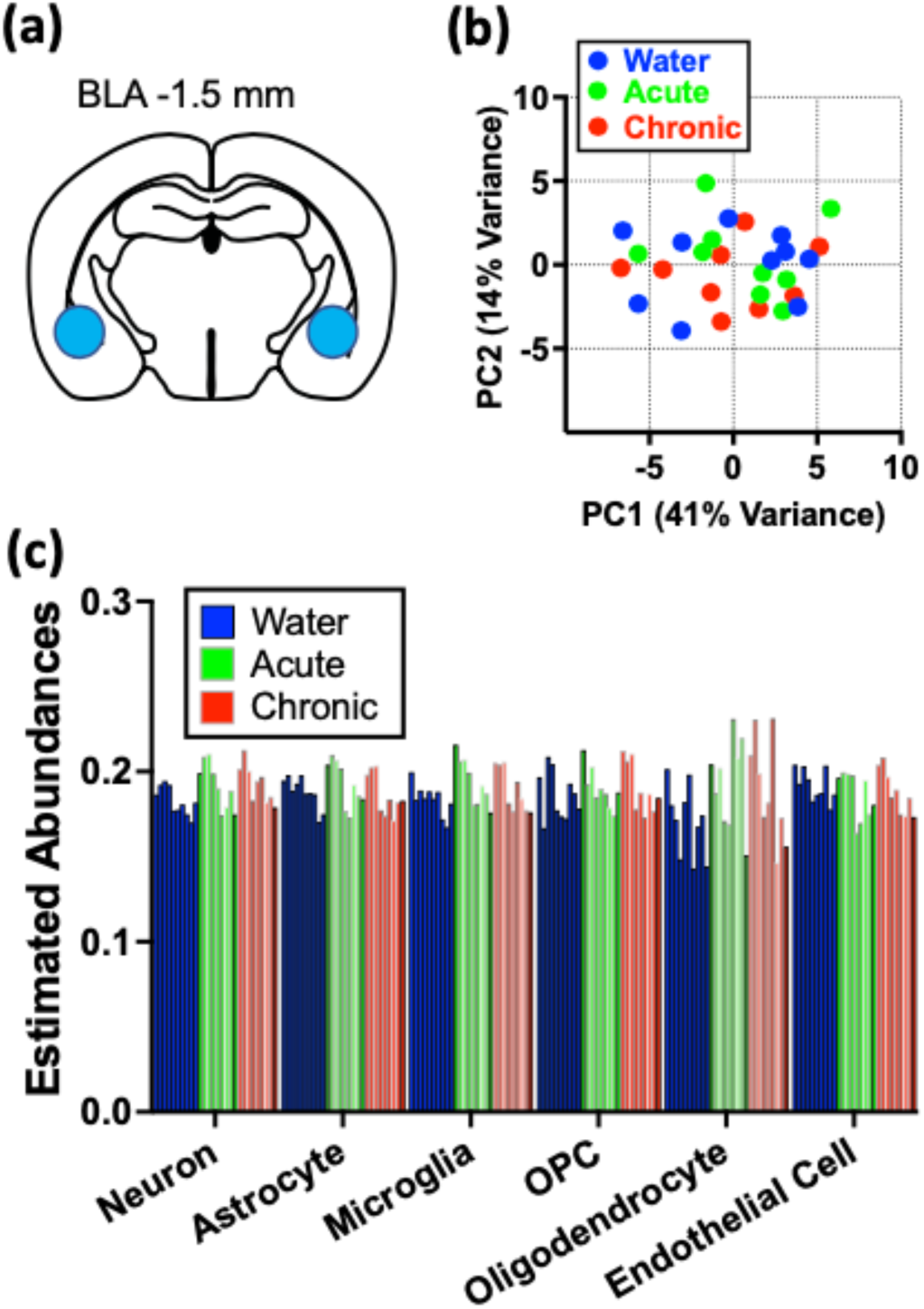
Initial assessment of sequencing results. (**a**) Diagram for amygdala tissue collection. (**b**) PCA plot showing no separation on alcohol drinking condition over the first two principal components, explaining 55% of the variation in total. (**c**) Bar graphs showing the estimated cell type abundance for seven relevant cell types as determined by cell type deconvolution analysis. Each bar represents a single sample.

To identify genes exhibiting significantly altered expression due to alcohol drinking, we next calculated the expression level of each transcript based on the number of transcripts per million reads, followed by normalization of reads. Initially, using the DESeq2 package [31] with parameters of p < 0.05 and |Log(fold change)| > 0, we identified 1300 and 1384 differentially expressed genes between Acute and Water drinking groups and between Chronic and Water drinking groups, respectively (Figure 3a and b). Then, we used the more stringent false discovery rate (FDR)-adjusted p-value cutoff of 0.05 to trim potential false positive results. We further identified 30 (Acute vs. Water, up-regulated: 23 and down-regulated: 6) and 97 (Chronic vs. Water, up-regulated: 36 and down-regulated: 61) differentially expressed genes (Figure 3c and Table 1, 2). These studies identified a number of robustly and significantly differentially expressed genes in the amygdala as a result of acute or chronic ethanol voluntary drinking.

**Figure 3.**
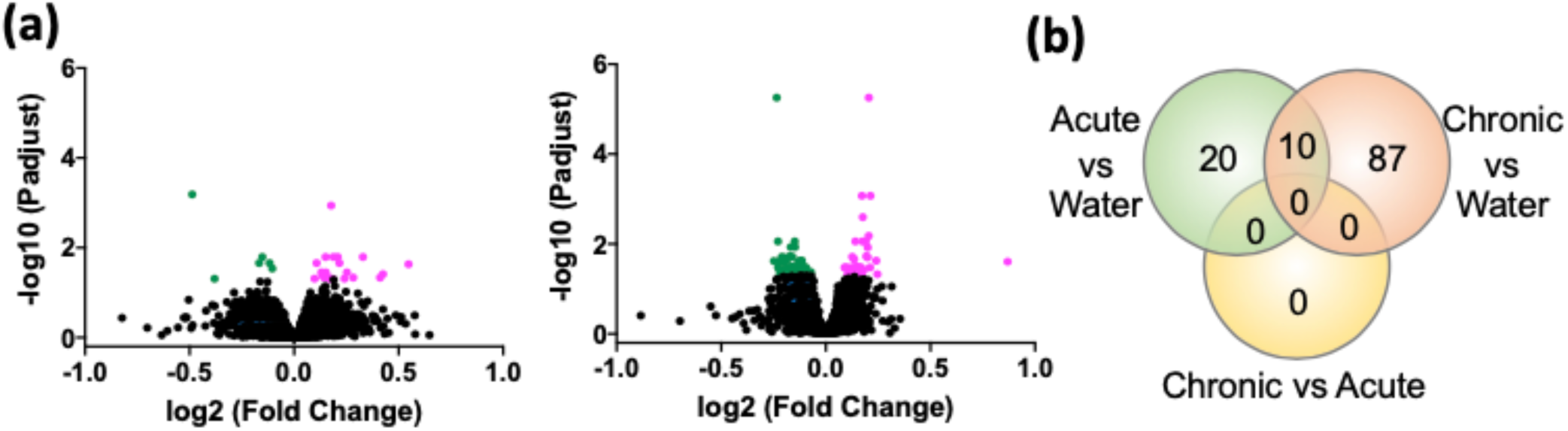
Differential gene expression analysis. (**a**) Volcano plot showing the fold change and adjusted p-value for each gene. Genes with significant up-regulation (p adjusted < 0.05) in Acute Drinking group (left) and Chronic Drinking group (right) are colored in magenta, and genes with significant down-regulation are colored in green. (**b**) Overlap Venn Diagrams of differentially expressed genes (DEGs) for comparisons across all drinking groups. P-adjusted value cutoff = 0.05.

**Table 1.**
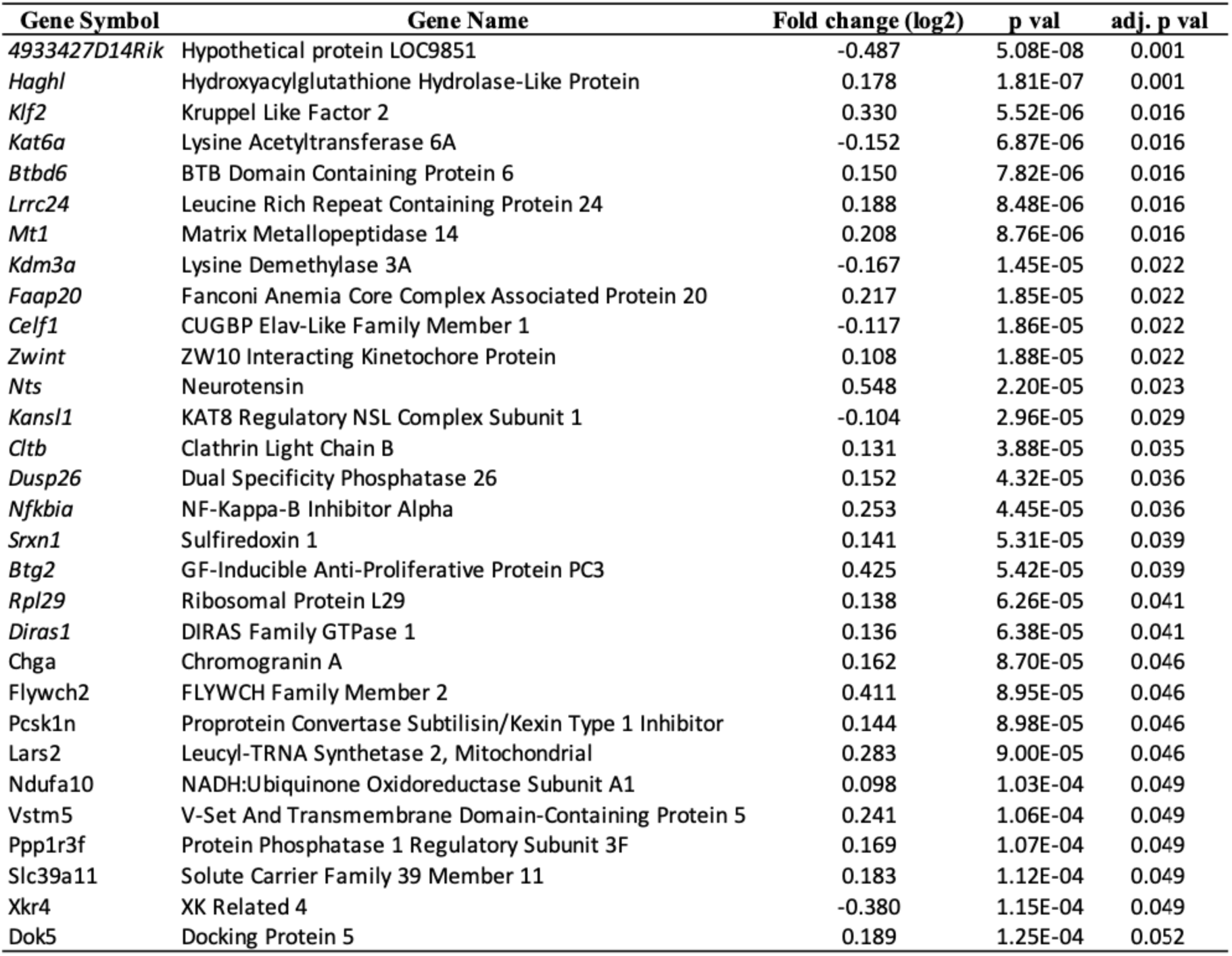
The 30 Acute Group-assocated DEGs.

**Table 2.**
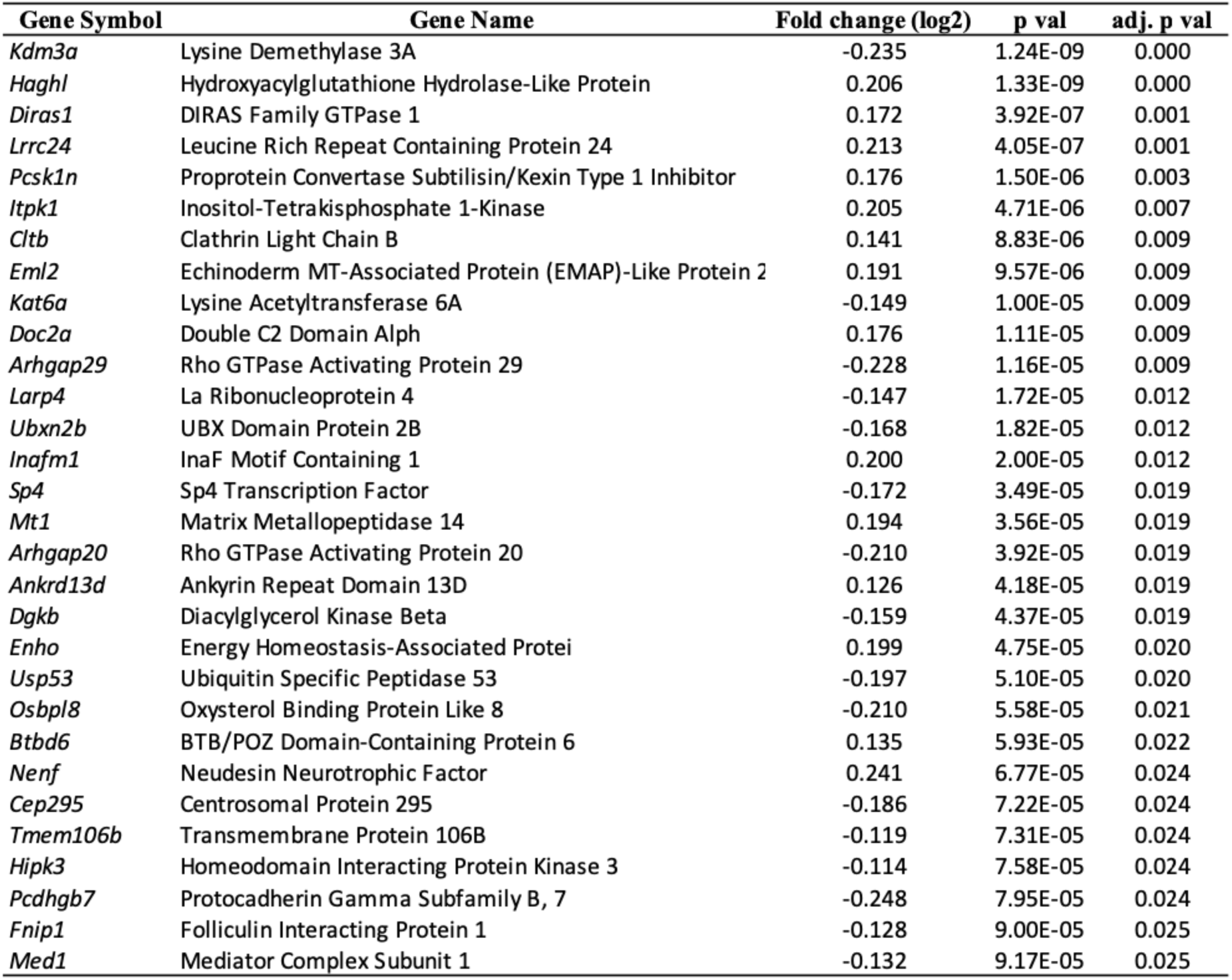
The 30 Chronic Group-assocated DEGs.

### GO and KEGG Gene Enrichment Analyses

To further identify networks of coordinately regulated genes that might point to alcohol-related specific biological functions, we next performed Gene Ontology (GO) and Kyoto Encyclopedia of Genes and Genomes (KEGG) Pathway enrichment analyses. We found that the primary effects of acute and chronic alcohol drinking were related to ribosome, cytoplasmic translation, chromatin binding, and histone modification pathways (Figure 4 and Figure 5). Interestingly, among those GO enrichment terms, “myelin sheath” (FDR-adjusted p = 0.0003) was in the top 5 up-regulated pathways, suggesting alcohol drinking affects molecular and cellular mechanisms underlying oligodendrocyte maturation and myelination, consistent with previous reports that demonstrated glial dysfunction in AUD pathophysiology [38]. These findings suggest that chronic alcohol drinking induces neuroadaptations mediated by glia-specific molecular alterations in the amygdala.

**Figure 4.**
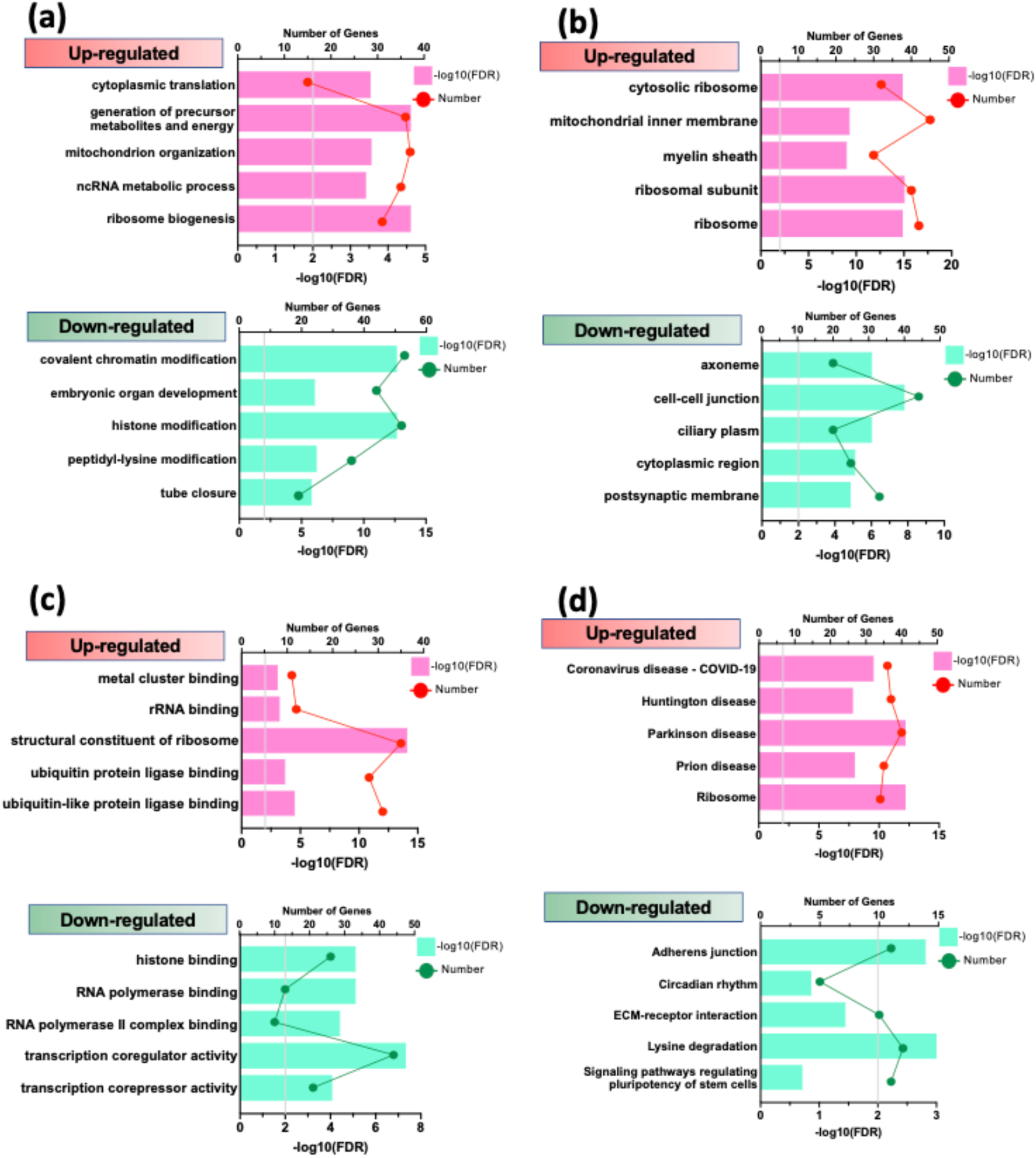
GO enrichment and KEGG pathway analysis of DEGs in response to Acute alcohol drinking. The ordinate represents the GO or KEGG terms, the upper abscissa indicates the number of genes in the GO/KEGG terms, and the lower abscissa indicates the level of significance of the enrichment (gray bar, FDR = 0.01) (**a**) Genes were categorized with the Biological Process domain. (**b**) Genes were categorized with the Cellular Component domain. (**c**) Genes were categorized with the Molecular Function domain. (**d**) The top 5 enriched pathways in the KEGG pathway analysis (gray bar, FDR = 0.01).

**Figure 5.**
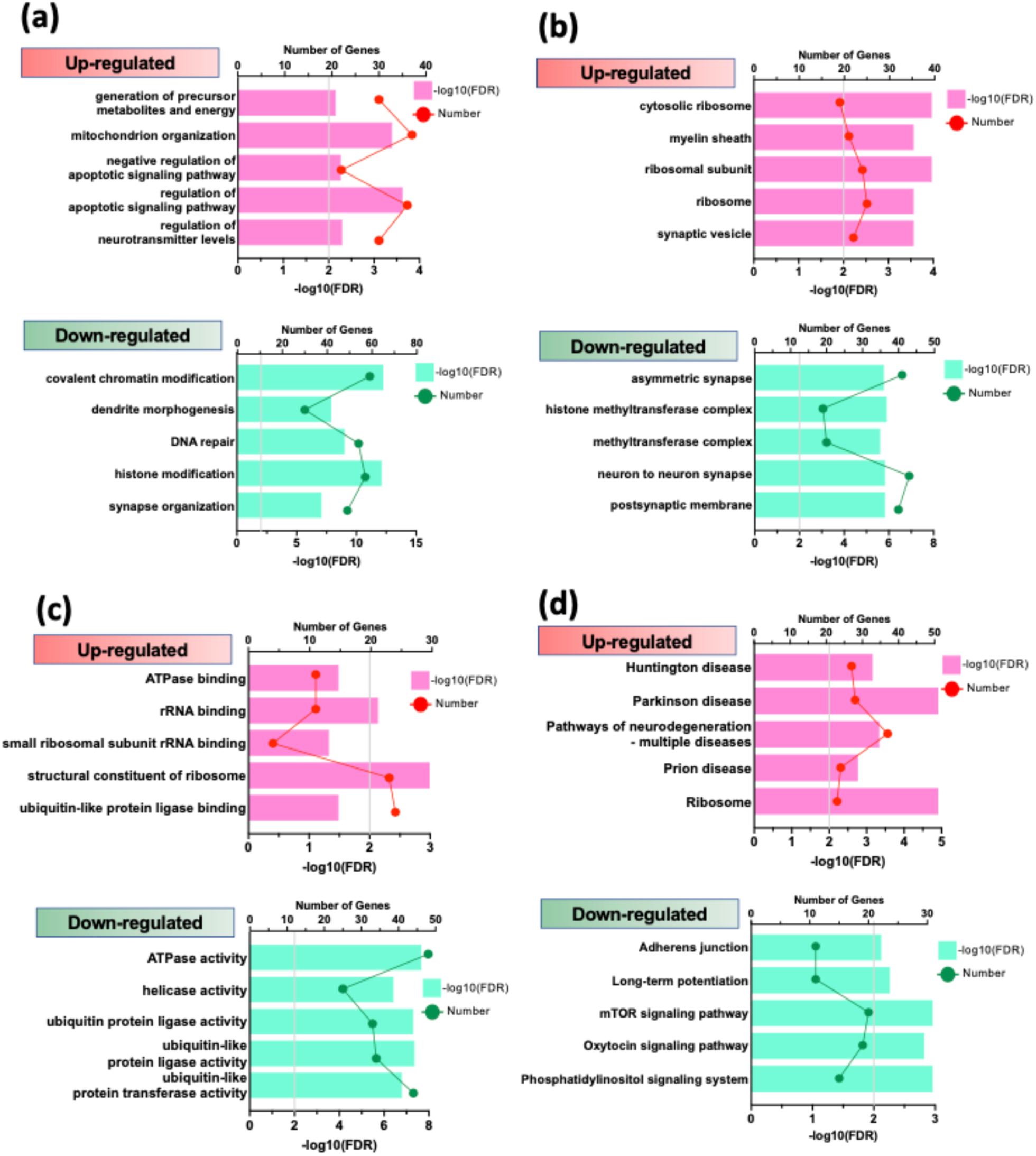
GO enrichment and KEGG pathway analysis of DEGs in response to Chronic alcohol drinking. The ordinate represents the GO or KEGG terms, the upper abscissa indicates the number of genes in the GO/KEGG terms, and the lower abscissa indicates the level of significance of the enrichment (gray bar, FDR = 0.01) (**a**) Genes were categorized with the Biological Process domain. (**b**) Genes were categorized with the Cellular Component domain. (**c**) Genes were categorized with the Molecular Function domain. (**d**) The top 5 enriched pathways in the KEGG pathway analysis (gray bar, FDR = 0.01).

### Identification of the potential regulatory pathways

To identify a list of hub genes, we next used STRING, a common online database for predicting protein-protein interaction networks. It revealed hub genes with 4-5 nodes, including histone deacetylase 2 (*Hdac2*), heterogeneous nuclear ribonucleoprotein M (*Hnrnpm*), histone deacetylase complex subunit sin3a (*Sin3a*), and chromodomain helicase DNA binding protein 1 (*Chd1*), particularly in Chronic Drinking condition (Figure 6). The *Hdac2, Sin3a* and *Chd1* are members of proteins associated with histone deacetylase activity. We then applied DEGs to the GeneGo MetaCore online database and identified 4 and 17 candidate upstream regulatory transcription factors in Acute and Chronic Drinking conditions, respectively (Table 3). SRY (sex determining region Y)-box transcription factor 17 (*Sox17*) stood out in both Acute and Chronic Drinking conditions. Since *Sox17* has been shown to regulate oligodendrocyte progenitor cell expansion and differentiation, this finding is consistent with our results from GO/KEGG pathway analyses, particularly “myelin sheath” (Figure 4 and 5). In addition, chronic drinking seems to drive multiple transcription factors including cAMP responsive element binding protein 1 (*Creb1*), which has been well-known to be involved in fear memory processing and alcohol exposure. Together, the findings suggest that voluntary alcohol consumption affects epigenetic changes via histone deacetylation pathways, and oligodendrocyte-related transcriptional factor, *Sox17*.

**Figure 6.**
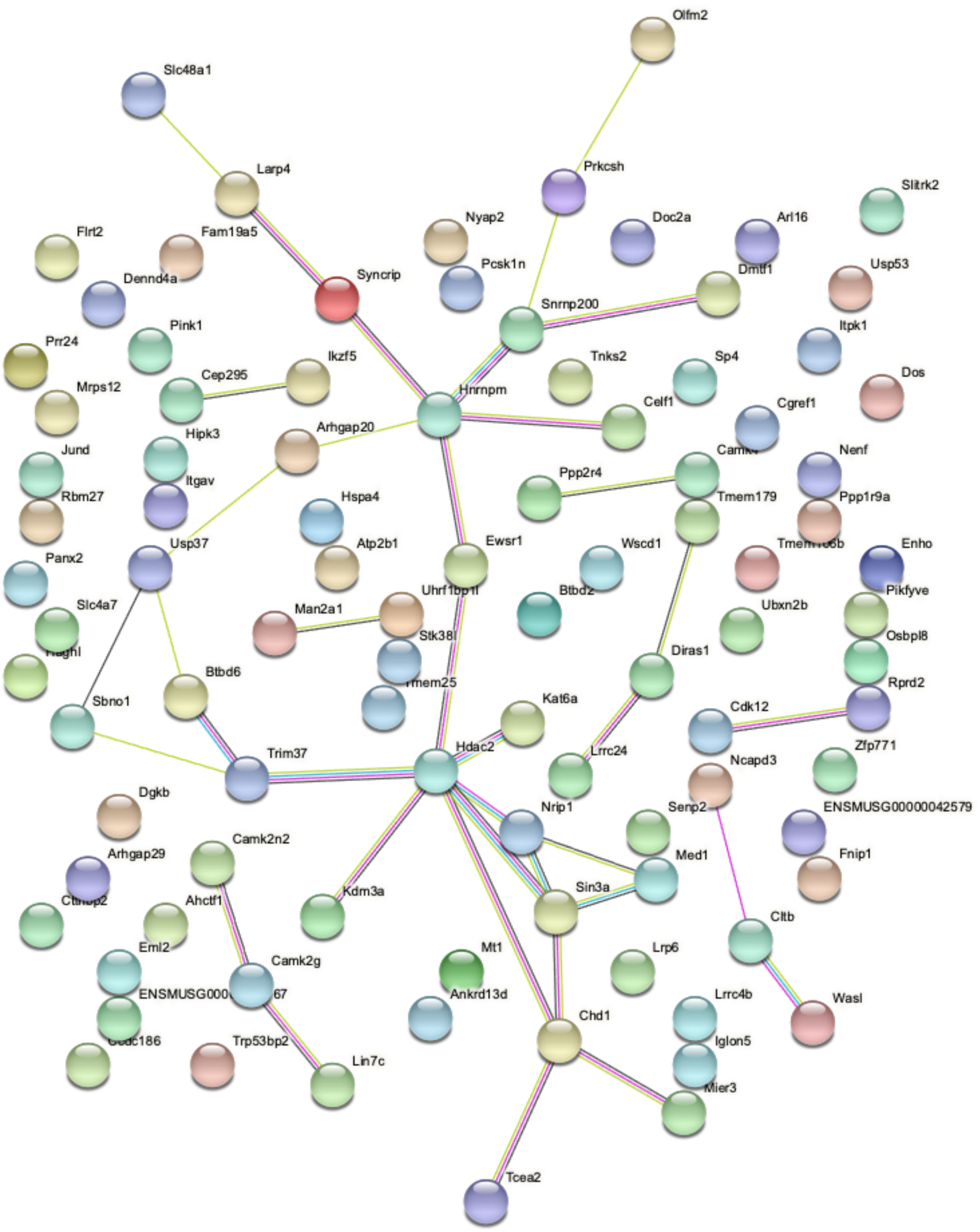
The protein-protein interaction (PPI) network analysis using STRING database. The 97 DEGs from the Chronic drinking group were input into STRING database and archived 95 nodes and 38 edges, with PPI enrichment p-value < 0.00352.

**Table 3.** Candidate Upstream Transcription Factors.

| Drinking Condition | Gene Symbol | Gene Name | Actual Targets | p-val | z-score |
| --- | --- | --- | --- | --- | --- |
| Acute | *Taf3* | TATA-Box Binding Protein Associated Factor 3 | 2 | 1.04E-04 | 16.02 |
|  | *c-Jun* | Jun Proto-Oncogene, AP-1 Transcription Factor Subunit | 7 | 2.94E-04 | 5.76 |
|  | *c-Fos* | Fos Proto-Oncogene, AP-1 Transcription Factor Subunit | 5 | 1.87E-04 | 5.27 |
|  | *Sox17* | SRY (Sex Determining Region Y)-Box Transcription Factor 17 | 22 | 3.60E-04 | 3.63 |
| Chronic | *Rbpj* | Recombination Signal Binding Protein for Immunoglobulin Kappa J Region | 58 | 2.27E-11 | 7.18 |
|  | *Sox17* | SRY (Sex Determining Region Y)-Box Transcription Factor 17 | 69 | 4.86E-11 | 6.70 |
|  | *Foxp3* | Forkhead Box P3 | 57 | 4.62E-08 | 5.69 |
|  | *Tal1* | TAL BHLH Transcription Factor 1, Erythroid Differentiation Factor | 61 | 9.83E-08 | 5.45 |
|  | *Creb1* | CAMP Responsive Element Binding Protein 1 | 44 | 1.29E-07 | 5.74 |
|  | *Ets1* | ETS Proto-Oncogene 1, Transcription Factor | 53 | 9.50E-07 | 5.09 |
|  | *c-Myc* | MYC Proto-Oncogene, BHLH Transcription Factor | 37 | 6.78E-05 | 4.22 |
|  | *Runx1* | Runt-Related Transcription Factor 1 | 46 | 1.17E-04 | 3.94 |
|  | *Gata-2* | GATA Binding Protein 2 | 23 | 1.63E-04 | 4.20 |
|  | *E2f1* | E2F Transcription Factor 1 | 37 | 1.79E-04 | 3.92 |
|  | *Zfx* | Zinc Finger Protein X-Linked | 23 | 2.20E-04 | 4.10 |
|  | *Gabp* | GA Binding Protein Transcription Factor Subunit Alpha | 35 | 3.97E-04 | 3.69 |
|  | *Cebpe* | CCAAT/Enhancer Binding Protein (C/EBP), Epsilon | 4 | 6.07E-04 | 5.85 |
|  | *Yy1* | YY1 Transcription Factor | 13 | 7.79E-04 | 3.95 |
|  | *Ash2* | ASH2 Like, Histone Lysine Methyltransferase Complex Subunit | 21 | 2.54E-03 | 3.21 |
|  | *Glis3* | GLIS Family Zinc Finger 3 | 20 | 2.82E-03 | 3.19 |
|  | *Klf9* | Kruppel Like Factor 9 | 3 | 3.47E-03 | 4.93 |

### GWAS catalog and DisGeNET

To determine if our DEGs (adjusted p < 0.05) are associated with AUD, we used online genome-wide association studies (GWAS) catalog database. We found that 5 of our DEGs, including ATPase plasma membrane calcium transporting 1 (*Atp2b1*), heat shock protein family A member 4 (*Hspa4*), strawberry notch homolog 1 (*Sbno1*), solute carrier family member 7 (*Slc4a7*), and UBX domain protein 2b (*Ubxn2b*), from the Chronic Drinking group have previously been identified in GWAS of AUD.

To further compare our DEGs with previously reported findings in AUD, we also took advantage of the publicly available databases DisGeNET. We found that nuclear factor kappa B subunit 1 (*Nfkb1*) and neurotensin (*Nts*) from Acute Drinking group have been previously linked to AUD. Similarly, beside 5 DEGs identified from GWAS, we found 5 more DEGs, including calcium/calmodulin dependent protein kinase IV (*Camk4*), energy homeostasis associated (*Enho*), *Hdac2*, LDL receptor related protein 6 (*Lrp6*), *Slc4a7*, and SLIT and NTRK like family member 2 (*Slitrk2*)), which were previously linked to AUD in the literature.

### Validation of DEGs by qPCR

Seven genes including brain cytoplasmic RNA 1 (*Bc1*), BTG anti-proliferation factor 2 (*Btg2*), hydroxyacylglutathione hydrolase like (*Haghl*), leucine rich repeat containing 24 (*Lrrc24*), neudesin neurotrophic factor (*Nenf*), and lysine demethylase 3A (*Kdm3a*) were used for qPCR analysis to validate the expression profiles obtained by bulk RNA-seq (Table 4). Consistent with the RNA-seq findings, in all cases, the relative fold change of gene expression was in the same direction in Acute and Chronic drinking groups.

**Table 4.** Validation of DEGs by qPCR.

|  |  | RNA-seq |  | qPCR |  |  |
| --- | --- | --- | --- | --- | --- | --- |
|  |  | Fold change (log2) |  | Fold change |  |  |
|  | Gene | Acute | Chronic | Water | Acute | Chronic |
| **up-regulated** | *Bc1* | 0.001 | 0.223 | 1.00 ± 0.11 | n.a. | 1.27 ± 0.13 |
|  | *Btg2* | 0.425 | 0.206 | 1.00 ± 0.09 | 1.40 ± 0.15* | n.a. |
|  | *Haghl* | 0.178 | 0.205 | 1.00 ± 0.10 | 1.35 ± 0.10* | 1.57 ± 0.14** |
|  | *Lrrc24* | 0.188 | 0.213 | 1.00 ± 0.13 | 1.37 ± 0.10* | 1.04 ± 0.12 |
|  | *Nenf* | 0.200 | 0.241 | 1.00 ± 0.09 | n.a. | 1.34 ± 0.10* |
|  | *Nts* | 0.548 | 0.239 | 1.00 ± 0.08 | 1.52 ± 0.18* | n.a. |
| **down-regulated** | *Kdm3a* | -0.167 | -0.235 | 1.00 ± 0.09 | 0.67 ± 0.07* | 0.71 ± 0.05* |

## Discussion

We have characterized the transcriptome level response to acute and chronic intermittent ethanol drinking with a 2-bottle choice drinking procedure, and identified sets of significant DEGs, distinct GO and KEGG pathways, hub genes, and upstream transcriptional factors that are sensitive to ethanol drinking in the mouse amygdala. Our results demonstrate that both acute and chronic ethanol drinking can impact on similar biological processes including translational machinery, epigenetic modifications, synaptic plasticity, and neurological disorders in the amygdala. Many of the genes identified in this mouse model of amygdala transcriptional regulation with alcohol had previously been associated with human AUD via GWAS or other genomic approaches, supporting the use of this model for translational, mechanistic studies. Furthermore, the findings also add to a body of evidence indicating that ethanol exposure leads to molecular and cellular alterations in non-neuronal cell types, such as oligodendrocytes.

Notably, we did not observe FDR-significant differences in DEGs and GO/KEGG pathways between Acute and Chronic drinking groups. A single bout of voluntary ethanol consumption resulted in molecular changes in the amygdala similar to those altered by repeated alcohol drinking, suggesting that acute ethanol drinking is sufficient to trigger critical molecular adaptations, possibly leading to future addiction with repeated alcohol exposure. As acute behavioral responses to alcohol have predictive value regarding risk for long-term alcohol drinking behavior in humans [39] and animal models [40], our results indicate striking overlapping regulation of amygdala-specific gene expression by alcohol regardless of the number of drinking episodes.

Many of the genes we identified were of great interest given prior findings. Notably, neurotensin (Nts) and its receptors have been implicated as contributing to the behavioral effects of alcohol in animal models. Chronic ethanol exposure increased Nts expression in the dorsal striatum [41], whereas ethanol decreased the expression of Nts receptors in both the nucleus accumbens (NAcc) and midbrain [42]. Furthermore, recent work has demonstrated that Nts-expressing neurons in the CeA contribute to the voluntary consumption of alcohol [43]. These findings are consistent with our results indicating an increase in Nts expression in Acute Drinking group, as the micropunches included the CeA in our samples. Interestingly, our recent studies demonstrated that the Nts receptor 2 (Ntsr2) is highly expressed in the BLA Thy1+ neurons that strongly project to the NAcc [44]. Therefore, our result provides a novel insight at a circuit level into how the amygdala subnuclei, CeA-Nts and BLA-Ntsr2, may interactively mediate voluntary and cue-induced alcohol drinking behaviors.

Our study revealed many other well-known pathways affected by alcohol drinking. Amongst these, first, translational machinery including rRNA binding, ribosome and ribosomal subunits seems to be positively affected by alcohol drinking. Consistent with previous studies from different brain areas [45], these results suggest that alcohol exposure also similarly affects translation of proteins in the amygdala, and ultimately leads to neuroadaptation via re-organization of synaptic structures, synaptic proteins and neurotransmitter receptors, such glutamate and GABA receptors [45]. Second, we found several enrichment terms related to chromatin remodeling, histone modification and DNA methylation in GO/KEGG analyses. Similarly, we also identified hub genes, mostly involved in histone deacetylation activity, including *HDAC2*. It was recently shown that there is increased HDAC2 level and activity in the amygdala of *P* rats, an alcohol-preferring rat line, and acute ethanol injection decreased HDAC2 activity and subsequently reduced voluntary ethanol intake [46]. These results suggest that alcohol drinking affects gene expression by potentially regulating epigenetic alterations, particularly histone modifications via HDAC2. Third, we found that pathways related to neurodegenerative diseases, including Huntington disease and Parkinson disease, are enriched in the KEGG analysis. Since the brain is a major target for the actions of alcohol, and heavy alcohol consumption has long been associated with brain damage as a risk factor [47], our study also confirms that alcohol consumption triggers similar molecular pathological pathways involved in neurodegenerative diseases. Fourth, one of interesting GO terms in our analysis is “myelin sheath.” Since our estimation of different cell types did not detect any discrepancy between samples from different drinking groups, the findings indicate that gene expression involved in myelin sheath formation and maintenance is affected by alcohol drinking. Interestingly, this pathway was up-regulated in both the Acute and Chronic Drinking groups, suggesting that a potential molecular recovery mechanism may be activated following myelination damages after alcohol drinking.

While genes are co-expressed forming functional networks, identifying upstream regulators of these genes and networks can provide insight into cellular function and lead to a potential therapeutic intervention. In our study, we observed that many of the DEGs from Acute and Chronic groups are regulated by *Sox17*. Since Sox17 regulates OPC proliferation and differentiation to oligodendrocytes via Wnt/β-catenin signaling pathway [48, 49], alcohol drinking seems to directly impact Sox17 transcription factor and OPC and consequently myelination in the amygdala. Notably, our group previously reported that amygdala-dependent Wnt/β-catenin signaling pathway is also involved in fear memory consolidation [50]. Together these data, along with the above myelin sheath findings in our pathway analyses, provides strong evidence for a role of oligodendrocyte alterations in the aftermath of ethanol consumption.

In summary, we identified alcohol-sensitive amygdala-associated candidate genes and pathways targeted by measuring genome-wide transcriptomic analyses. Consistent with previous gene expression studies, we found that voluntary alcohol consumption, regardless the number of drinking episodes, results in similar gene expression changes in ribosome-related/translational pathways, myelination, chromatin-binding, and histone modification. These genes and pathways suggest convergence of human GWAS and molecular studies with amygdala transcription data from mouse drinking models. Future studies will use advanced cell targeting techniques to validate the roles of identified genes in neural adaptation processes mediating the progression from acute to chronic alcohol intake.

## Notes

### Competing Interest Statement

The authors have declared no competing interest.

